# Sorbitol uptake and oxygen transfer shape *AOX1* promoter induction under methanol-free conditions in *Komagataella phaffii* lacking formate dehydrogenase

**DOI:** 10.1101/2025.04.01.646572

**Authors:** Cristina Bustos, Rocio Cozmar, Julio Berrios, Patrick Fickers

## Abstract

For decades, expression systems based on the methanol-regulated *AOX1* promoter (p*AOX1*) from the alcohol oxidase 1 gene have served as a benchmark for recombinant protein (rProt) production in *Komagataella phaffii*. However, methanol-free processes are increasingly being developed to overcome the drawbacks of methanol utilization, particularly its toxicity and flammability. The use of formate as a p*AOX1* inducer in combination with sorbitol, a non-repressive carbon source, has emerged as a promising alternative to methanol-based expression systems. Recently, we demonstrated that formate derived from the tetrahydrofolate-mediated one-carbon (THF-C1) metabolism accumulates in *K. phaffii* cells deficient in formate dehydrogenase (FdhKO) when grown in sorbitol-based methanol-free medium. Using the lipase CalB from *Candida antarctica* as a model protein, we observed that rProt productivity in an FdhKO strain grown on sorbitol was comparable to that of an Fdh-proficient strain grown on methanol. However, sorbitol is inefficiently metabolized in *K. phaffii*, leading to a low growth rate and potentially limiting rProt productivity due to insufficient energy and carbon supply.

Here, we increased sorbitol uptake rate, and thus improved sorbitol metabolism, by overexpressing the gene encoding sorbitol dehydrogenase (*SOR1*) in an FdhKO strain. Our results demonstrate that while increased sorbitol metabolism promotes biomass formation, it reduces p*AOX1* induction, as evidenced by lower formate accumulation and decreased rProt productivity, both for intracellular eGFP and secreted proteins namely CalB lipase and glucose oxidase GOx from *Aspergillus niger* in *SOR1*-overexpressing strains. Additionally, oxygen availability for cells influences these dynamics, with lower oxygen transfer favoring higher p*AOX1* induction due to increased formate accumulation in an FdhKO strain. Our data also suggests that at low oxygen transfer and low sorbitol uptake rate, the proportion of cells in an induced state increased significantly. This work provides valuable insights into the interplay between sorbitol metabolism and oxygen transfer conditions, contributing to the development of improved recombinant protein production strategies in *K. phaffii*.

## Introduction

The methylotrophic yeast *Komagataella phaffii* (formerly *Pichia pastoris*) is widely used as a cell factory for recombinant protein (rProt) production (Ergün *et al*., 2021). For this purpose, the expression system based on the promoter from the alcohol oxidase 1 gene (p*AOX1*) has been a benchmark for decades. While p*AOX1* is tightly repressed in the presence of glucose or glycerol, it is strongly induced by methanol under non-repressive conditions. This allows the uncoupling of the biomass production phase on glycerol from the protein production phase using methanol as both a carbon source and inducer. However, methanol is toxic to cells and highly flammable, posing significant challenges for large-scale production processes. To address these limitations, methanol-free expression systems have been developed through promoter and strain engineering or by utilizing alternative inducers for p*AOX1* in combination with a non-repressive carbon source (for a review, see Singh and Narang, 2020; Ergün et al., 2021).

Sorbitol is a six-carbon polyol that does not repress p*AOX1* induction and can, therefore, be used as a co-substrate during methanol induction (Niu *et al*., 2013; Ergün *et al*., 2021). After being transported across the plasma membrane by a hexose transporter (Jordan *et al*., 2016), sorbitol is converted into fructose by sorbitol dehydrogenase. Fructose is subsequently phosphorylated by hexokinase before entering the glycolysis pathway (Figure S1). The *K. phaffii* growth rate on sorbitol is low (0.03 h^−1^; Niu et al., 2013; Singh and Narang, 2023), with a substrate to biomass conversion yield (Y_x/s_) of 0.56 g g^-1^ (Singh and Narang, 2023). Recently, formate has emerged as a promising alternative inducer of p*AOX1*-driven rProt production (Tyurin and Kozlov, 2015; Jayachandran *et al*., 2017; Singh and Narang, 2020). In the methanol dissimilation pathway (Berrios *et al*., 2022), formate is generated from formaldehyde by formaldehyde dehydrogenase (Fld) and further converted into carbon dioxide by formate dehydrogenase (Fdh) (Hartner and Glieder, 2006); Figure S2). Additionally, formate is an intermediate in the tetrahydrofolate-mediated one-carbon (THF-C1) metabolism, which plays a role in several anabolic pathways, including *de novo* purine synthesis (Christensen and Mackenzie, 2006). Recently, we demonstrated that formate from THF-C1 metabolism accumulates in *K. phaffii* cells deficient in formate dehydrogenase (FdhKO) when grown in a sorbitol-based methanol-free medium. This endogenous formate was shown to be sufficient to trigger p*AOX1* induction without the need for external inducer supplementation. This finding is particularly significant because any p*AOX1*-based expression system can be converted into a self-induced system in an FdhKO strain. When cells are cultivated in a mixture of glycerol and sorbitol, rProt synthesis is initiated at the end of the exponential growth phase, upon glycerol depletion from the culture medium. Using the secreted lipase CalB from *Candida antarctica* as a model protein, comparable productivities were obtained with an FdhKO CalB strain grown on sorbitol and an Fdh-expressing CalB strain grown on methanol (Bustos *et al*., 2024).

In *K. phaffii*, energy generation is a known bottleneck for rProt synthesis in methanol-free p*AOX1*-based processes (Feng *et al*., 2022). Since the low sorbitol consumption rate by *K. phaffii* could potentially limit rProt productivity in an FdhKO strain, we aimed to enhance sorbitol metabolism in an FdhKO strain and assess the impact on rProt productivity. As oxygen availability to cells can also be a limiting factor for rProt synthesis, this parameter was also considered for strains optimized for sorbitol metabolism. Our results highlight that the combination of poor sorbitol utilization and low oxygen transfer condition enhances formate accumulation in *K. phaffii* FdhKO cells, thereby promoting higher rProt productivity.

## Experimental procedures

### Strains and media and culture conditions

The *Escherichia coli* and *K. phaffii* strains used in this study are listed in Table 1 and S1. *E. coli* was grown at 37°C in Luria-Bertani medium (LB), supplemented with antibiotics as follows: 100 µg mL^-1^ ampicillin, 50 µg mL^-1^ kanamycin, or 25 µg mL^-1^ zeocin. *K. phaffii* was grown at 30 °C either on YPD medium (20 g L^-1^ glucose, 10 g L^-1^ Difco yeast extract, and 10g L^-1^ Difco bacto peptone) or in YNB medium (1.7 g L^-1^ Difco YNB w/o ammonium chloride and amino acids, 5 g L^-1^ NH_4_Cl and, 0.4 mg L^-1^ biotin, 100 mM potassium phosphate buffer, pH 6.0) supplemented with 10 g L^-1^ sorbitol (YNBS), 1.75 g L^-1^ glycerol (YNBG0), 1.75 g L^-1^ glycerol and 1.75 g L^-1^ sorbitol (YNBGS2), 1.75 g L^-1^ glycerol and 3.5 g L^-1^ sorbitol (YNBGS4) or 20 g L^-1^ sorbitol (YNBS2). *K. Phaffii* transformants were selected on YPD agar plates, supplemented with 25 µg mL^-1^ zeocin (YPD-Zeo). Precultures were operated as previously described (Bustos *et al*., 2024). Cultures were performed in Erlenmeyer shake flasks (50 mL or 250 mL) or in microbioreactors (BioLector 2, m2p-labs, Baesweiler, Germany) as previously described (Bustos et al. 2024). Cultures under different oxygen transfer conditions (OTC) were carried out as previously described in 250 mL Erlenmeyer flasks containing varying medium volumes to benchmark oxygen transfer coefficients (K_L_a) of 10, 50, and 100 h^-1^ (Gorczyca *et al*., 2020).

**TABLE 1.** *K. phaffii* strains used in this study.

| Number | Names | Parental strain, genotype | Reference |
| --- | --- | --- | --- |
| RIY232 | GS115pro | GS115, *HIS4* | (Theron *et al.*, 2019) |
| RIY230 | Fdh eGfp | GS115, pAOX1-eGFP | (Velastegui *et al.*, 2019) |
| RIY308 | Fdh CalB | GS115, pAOX1-αMF-CalB | (Velastegui *et al.*, 2019) |
| RIY536 | FdhKO eGfp Z<sup>+</sup> | RIY230, *fdh1Δ*, pAOX1-eGFP Zeo<sup>+</sup> | (Bustos *et al.*, 2024) |
| RIY537 | FdhKO CalB Z<sup>+</sup> | RIY308, *fdh1Δ*, pAOX1-αMF-CalB Zeo<sup>+</sup> | (Bustos *et al.*, 2024) |
| RIY540 | FdhKO eGfp | RIY536, *fdh1Δ*, pAOX1-eGFP Zeo<sup>-</sup> | (Bustos *et al.*, 2024) |
| RIY561 | FdhKO CalB | RIY537, *fdh1Δ*, pAOX1-αMF-CalB Zeo<sup>-</sup> | (Bustos *et al.*, 2024) |
| RIY529 | SdOE | RIY232, pGAP-SOR1 | This work |
| RIY530 | StOE | RIY232, pGAP-PpHXT1 | This work |
| RIY532 | St&SdOE | RIY232, pGAP-SOR1, pGAP-PpHXT1 | This work |
| RIY643 | FdhKO-SdOE eGfp | RIY540, *fdh1Δ*, pAOX1-eGFP, pGAP-SOR1 | This work |
| RIY562 | FdhKO-SdOE CalB | RIY561, *fdh1Δ*, pAOX1-αMF-CalB, pGAP-SOR1 | This work |
| RIY655 | FdhKO GOx | *fdh1Δ*, pAOX1-αMF-Gox | Lab stock (unpublished) |
| RIY656 | FdhKO SdOE GOx | *fdh1Δ*, pAOX1-αMF-GOx, pGAP-SOR1 | Lab stock (unpublished) |

### Construction of plasmid and K. phaffii strains

The general genetic techniques were as described elsewhere (Bustos *et al*., 2024). The plasmids and primers used are listed in Table S1 and S2. The SdOE and StOE expression vectors were constructed using the GoldenPiCS system (Prielhofer *et al*., 2017). The gene PAS_chr1-1_0490 (*SOR1*) was PCR-amplified using *K. Phaffii* GS115pro genomic DNA as a template, and an internal BpiI recognition sequence in gene PAS_chr1-1_0490 (*SOR1*) was removed by overlapping PCR with primers SOR1-Fw/ SOR1-Rv. In the gene PAS_chr1-4_0570 (*PpHXT1*), internal BpiI and BsaI recognition sequences were removed through in silico design by selecting alternative commonly used synonymous codons, and the synthetic gene was designed with the addition of external BsaI recognition sequences. The corresponding synthetic DNA fragment was obtained from Eurofin Genomic (Ebersberg, Germany). The resulting PCR product and synthetic gene fragment were cloned into plasmid A2 (BB1-23) at the BsaI restriction site, yielding plasmids RIE341 (A2_BB1_23_*SOR1*) and RIE343 (A2_BB1_23_Pp*HXT1*), respectively. Positive constructs were verified by colony and plasmid PCR with M13-Fw/M13-Rv. Plasmid RIE354 (p*GAP*-*SOR1*-ScCYC1tt) and RIE355 (p*GAP*-Pp*HXT1*-ScCYC1tt) were assembled by Golden Gate assembly from the plasmids A4 (BB1_12_p*GAP*), C1 (BB1_34_ScCYC1tt), D12 (BB3aZ_14), with RIE341 and RIE343, respectively, using BpiI as the restriction enzyme. The correctness of the assembly constructs was verified by colony and plasmid PCR with primers pair pGAPInt-Fw/ScCYC1tt.Int-Rv.

The Loxp-Zeo-Loxp was released from plasmid D12 (BB3aZ_14) by HindIII and PstI restriction and cloned at the corresponding site of C12(BB3aK_AC) to yield plasmid RIE370 (BB3-AC-LoxP-Zeo-loxP). The double overexpression of *SOR1* and Pp*HXT1* was performed in three steps. First, plasmid RIE372 (BB2_AB-p*GAP*-*SOR1*-*SsCy1*tt) was obtained by assembly of parts from plasmid RIE341, A4, C1, and D4 using BsaI as the restriction enzyme. Second, plasmid RIP374 (BB2_BC1-P*GAP*-Pp*HXT1-RPS3*tt) was obtained by assembly of parts from RIE378, A4, C9 and D5 using BsaI. Finally, plasmid RIP378 (BB3a-p*GAP*-*SOR1*-SsCy1tt, p*GAP*-Pp*HXT1*-RPS3tt) was obtained by assembly of parts from RIE372, RIE374, and RIE370 were ensemble with BpiI. Plasmid construct confirmation was performed with colony and plasmid PCR with the couple of primers pGAP.Int-Fw/ScCYC1tt.Int-Rv, and pGAP.Int-Fw/ Rps3tt.Int-Rv respectively.

The construction of yeast strains RIY230 (Fdh eGfp), RIY308 (Fdh CalB), RIY536 (FdhKO eGfp Z^+^), RIY537 (FdhKO CalB Z^+^), RIY540 (FdhKO eGfp) and RIY561 (FdhKO CalB) is described elsewhere (Velastegui *et al*., 2019; Bustos *et al*., 2024). Strains RIY529 (p*GAP*-*SOR1*-SsCy1tt, hereafter SdOE strain), RIY530 (p*GAP*-Pp*HXT1*-SsCy1tt, hereafter StOE strain) and RIY532(p*GAP*-*SOR1*-SsCy1tt, p*GAP*-Pp*HXT1*-SsCy1tt, hereafter St&SdOE strain) were obtained by transformation of strain RIY232 with AscI linearized plasmids RIP354 (p*GAP*-Pp*HXT1*-SsCy1tt), RIP355 (p*GAP*-*SOR1*-SsCy1tt) and RIP378 (p*GAP*-*SOR1*-SsCy1tt, p*GAP*-Pp*HXT1*-RPS3tt), respectively. Overexpression of *SOR1* in the RIY540 (FdhKO eGfp) and RIY561 (FdhKO CalB) strains were performed by transformation with AscI linearized plasmid RIP355 (BB3-p*GAP*-*SOR1*-*SsCy1*tt) to yield strains RIY643 (*fdh1*Δ, p*AOX1*-e*GFP*, p*GAP*-*SOR1*, hereafter FdhKO-SdOE eGfp strain) and RIY562 (*fdh1*Δ, p*AOX1*-αMF-*CalB*, p*GAP*-*SOR1*, hereafter FdhKO-SdOE CalB strain). The genotype of RIY529, RIY530, RIY532, RIY643, and RIY564 strains was verified by PCR using the primers pGAP.Int-Fw/ScCYC1tt.Int-Rv, pGAP.Int-Fw/ Rps3tt.Int-Rv, which annealed to the p*GAP*, ScCYC1tt, and Rps3ttregions, respectively. A schematic representation of the strain genotype is presented in Figure S5.

### Analytical methods

Cell growth was monitored either by optical density at 600nm (OD_600_) or dry cell weight (DCW). An OD_600_ value of 1 was found to correspond to 0.65 gDCW L^-1^. Sorbitol and glycerol concentrations were determined using high-performance liquid chromatography as described elsewhere (Bustos *et al*., 2024).

Intracellular eGFP fluorescence was quantified using a BD Accuri C6 Flow Cytometer (BD Biosciences, San Jose, CA, USA), as described elsewhere (Sassi *et al*., 2016; Bustos *et al*., 2024). To calculate the total value of fluorescence in the cell population, the FL1-A median value (i.e., the eGFP fluorescence) was multiplied by the fraction of cells with eGFP fluorescence (i.e., induced cells). It was expressed in total fluorescence unit (TFU). During cultures in microbioreactor, the eGFP fluorescence was quantified at 620 nm (excitation at 488) and expressed as specific fluorescence (sFU). For that purpose, fluorescence values were divided by the biomass values. The specific fluorescence value of the parental strain GS115pro was subtracted from the specific fluorescence of eGFP-producing strains to normalize the data.

Formate concentration in the culture supernatant were measured using the formic acid assay kit (Megazyme Inc., Bray, Ireland) according to the manufacturer instructions. Lipase activity in the culture supernatant was determined by monitoring the hydrolysis of p-nitrophenylbutyrate (p-NPB), following the protocols detailed elsewhere (Fickers *et al*., 2003). Glucose oxidase activity was monitored using a glucose oxidase assay kit (Megazyme Inc., Bray, Ireland) according to the manufacturer instructions. For formate and enzymatic assay absorbance at specific wavelength were monitored over time using a SpectraMax M2 (Molecular Devices, San Jose, CA, USA). All assays were conducted in triplicate

## Results and Discussion

### Overexpression of the sorbitol dehydrogenase encoding gene increases the sorbitol uptake rate

In *K. phaffii*, low energy generation is known as a limitation for rProt production in methanol-free processes based on the p*AOX1* (Feng *et al*., 2022). During the rProt production phase, sorbitol is one of the most widely used co-substrates (Çelik *et al*., 2009; Niu *et al*., 2013; Carly *et al*., 2016). Unfortunately, the ability of *K. phaffii* to catabolize sorbitol is low, potentially limiting the energy and carbon provision required for rProt synthesis. Sorbitol transport across the plasma membrane and its conversion to fructose by sorbitol dehydrogenase (Figure S1) have been identified as potential bottlenecks in sorbitol assimilation (Akentyev *et al*., 2023). The genes encoding these proteins have not yet been reported in *K. phaffii*. In *Saccharomyces cerevisiae*, the hexose transporter Hxt15 has been identified as a versatile polyol transporter, including for sorbitol (Jordan *et al*., 2016). Additionally, the gene encoding sorbitol dehydrogenase (Sdh, *SOR1*) has been characterized in both *S. cerevisiae* and *Komagataella kurtzmanii* (Toivari *et al*., 2004; Akentyev *et al*., 2023). A protein BLAST search using the translated *SOR1* and *HXT15* sequences from *S. cerevisiae* as queries highlighted the genes PAS_chr1-1_0490 (*SOR1*, 55.6% identity) and PAS_chr1-4_0570 (*PpHXT1*, 51% identity) as putative Sdh and Hxt candidates, respectively (Figure S3 and S4).

To increase the sorbitol uptake rate, these two genes were expressed under the control of the constitutive p*GAP* promoter, either individually or in combination, in the GS115pro strain, a prototrophic derivative of strain GS115 (Table 1). In the resulting StOE strain (p*GAP-PpHXT1*), the sorbitol uptake rate and cell growth rate increased by 1.7 and 1.2-fold, respectively, compared to the parental GS115pro strain (Table 2). For the SdOE strain (p*GAP-SOR1*), these rates were enhanced by 5.0 and 3.3-fold, respectively. However, for the St&SdOE strain, which co-expressed both genes (p*GAP-SOR1*, p*GAP-PpHXT1*), the sorbitol uptake rate and cell growth rate were not further enhanced compared to the SdOE strain (1.06 and 1.03-fold, respectively). These findings demonstrate that the expression of the *SOR1* gene alone can overcome the limiting step of sorbitol assimilation in *K. phaffii*.

**TABLE 2.** Effects of *SOR1* and *PpHXT1* genes overexpression on specific growth rate and the specific consumption rate of sorbitol. Abbreviations: µ, specific growth rate; DCW, dry cell weight; q_s_, specific sorbitol uptake rate. These cultures were performed in triplicate 250 mL flasks containing 25 mL of medium. Standard deviations were less than 10% of the presented mean values.

| Strains (genotypes) | $\mu$<br>(h<sup>-1</sup>) | $q_s$<br>(g gDCW<sup>-1</sup> h<sup>-1</sup>) |
| --- | --- | --- |
| GS115pro (GS115 prototroph) | 0.036 | 0.03 |
| StOE (pGAP-PpHXT1) | 0.042 | 0.05 |
| SdOE (pGAP-SOR1) | 0.119 | 0.15 |
| St&SdOE (pGAP-SOR1, pGAP-PpHXT1) | 0.122 | 0.16 |

Abbreviations: $\mu$ , specific growth rate; DCW, dry cell weight; $q_s$ , specific sorbitol uptake rate. These cultures were performed in triplicate 250 mL flasks containing 25 mL of medium. Standard deviations were less than 10% of the presented mean values.

### Enhancing sorbitol metabolism decreases pAOX1 induction in FdhKO strains on methanol-free medium

In many rProt production processes, glycerol is commonly used as a carbon source for biomass formation, while a p*AOX1* non-repressive carbon source, such as sorbitol, is preferred during the rProt synthesis phase (Ergün *et al*., 2022). Therefore, biomass and eGFP fluorescence were monitored for strains FdhKO eGfp (*fdh1*Δ, p*AOX1-eGFP*) and FdhKO SdOE eGfp (*fdh1*Δ, p*AOX1-eGFP*, p*GAP-SOR1*) grown in microbioreactors in media containing glycerol and sorbitol in different ratios (1:0, 1:1, 1:2; YNBG0, YNBGS2, and NBGS4 medium, respectively. Figure 1). Both strains exhibited diauxic growth, consuming glycerol within the first 8 hours of culture, followed by a sorbitol consumption phase (Figure S6). While both strains grew similarly on glycerol, marked differences were observed during the sorbitol consumption phase for the FdhKO eGfp and FdhKO SdOE eGfp strains (e.g., 0.035 ± 0.001 and 0.129 ± 0.001 h^−^1 on YNBGS4 medium, respectively). This confirms that the overexpression of *SOR1* increases the carbon assimilation and energy generation in *K. phaffii* grown on sorbitol.

**Figure 1.**
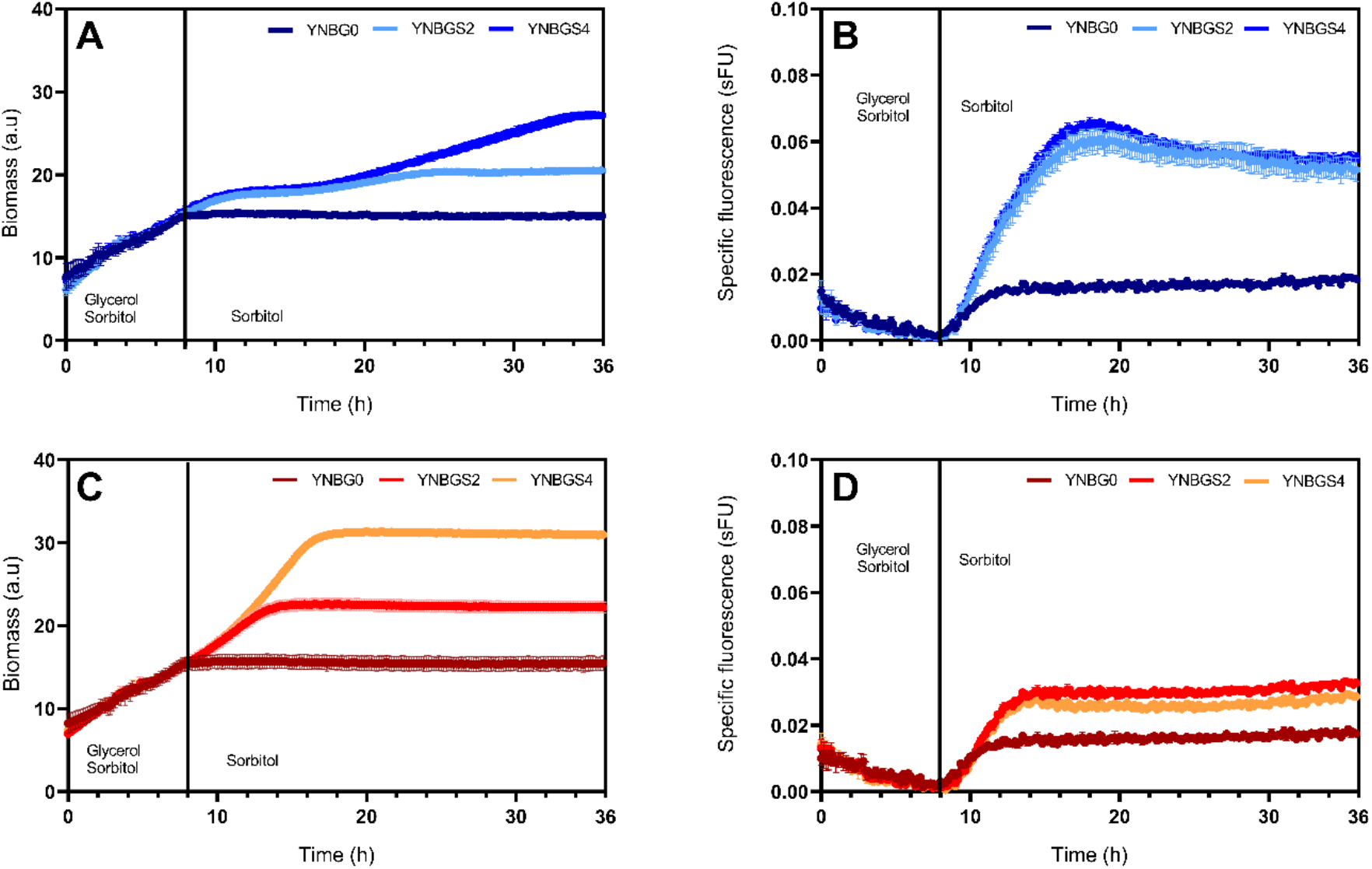
Biomass (panels A and C) and specific fluorescence (panels B and D) during the growth of FdhKO eGfp strain (blue colors) and FdhKO SdOE eGfp strain (orange colors) in YNB minimal medium containing different mixtures of glycerol and sorbitol (1:0, 1:1, 1:2; medium YNBG0, YNBGS2, and YNBGS4, respectively). Cells were grown in a BioLector system, and data represent the mean and standard deviation of triplicate cultures. sFU: specific fluorescence unit; a.u: arbitrary units.

Upon glycerol depletion in YNBG0 medium (in the absence of sorbitol), the specific fluorescence signals increased slightly within the same range for both the FdhKO eGfp and FdhKO SdOE eGfp strains (i.e., between 0.015 and 0.014 sFU), likely due to the derepression of p*AOX1* following glycerol depletion. Conversely, in sorbitol-containing media (YNBGS2 and YNBGS4), eGFP-specific fluorescence markedly increased for the FdhKO eGFP strain, reaching maximal values of 0.060 and 0.065 sFU, respectively. In contrast, the increase was less pronounced in the FdhKO SdOE eGfp strain, with maximal values of 0.032 and 0.028 sFU, respectively. These findings highlight that p*AOX1* induction is lower at higher sorbitol uptake rates (i.e. upon *SOR1* expression) and that sorbitol concentration does not significantly impact the strength of p*AOX1* induction.

Similar experiments were conducted using two secreted enzymes, lipase B (CalB) from *Candida antarctica* and glucose oxidase (GOx) from *Aspregillus niger*, as model proteins. Cell growth and respective enzyme activities, namely lipase or glucose oxidase, were quantified for strains FdhKO CalB (*fdh1*Δ, p*AOX1-αMF-CALB*), FdhKO SdOE CalB (*fdh1*Δ, p*AOX1-αMF-CALB*, p*GAP-SOR1*), FdhKO GOx (*fdh1*Δ, p*AOX1-αMF-GOX*) and FdhKO SdOE GOx (*fdh1*Δ, p*AOX1-αMF-GOX*, p*GAP-SOR1*) grown on sorbitol in shake flasks (Figure 2). After 36 hours of culture, both FdhKO SdOE strains showed higher biomass compared to their non-*SOR1*-expressing counterparts, with increases of 4.8 and 6.7-fold observed for the CalB and GOx expressing strains, respectively (Figure 2A, C). In contrast, these strains exhibited a 3.5 and 3.6-fold decrease in specific lipase and glucose oxidase activities, respectively, compared to their FdhKO counterparts (Figure 2B, D). These results for the secreted proteins align with the trend observed in the eGFP strains (i.e., FdhKO eGfp and FdhKO SdOE eGfp strains, Figure 1).

**Figure 2.**
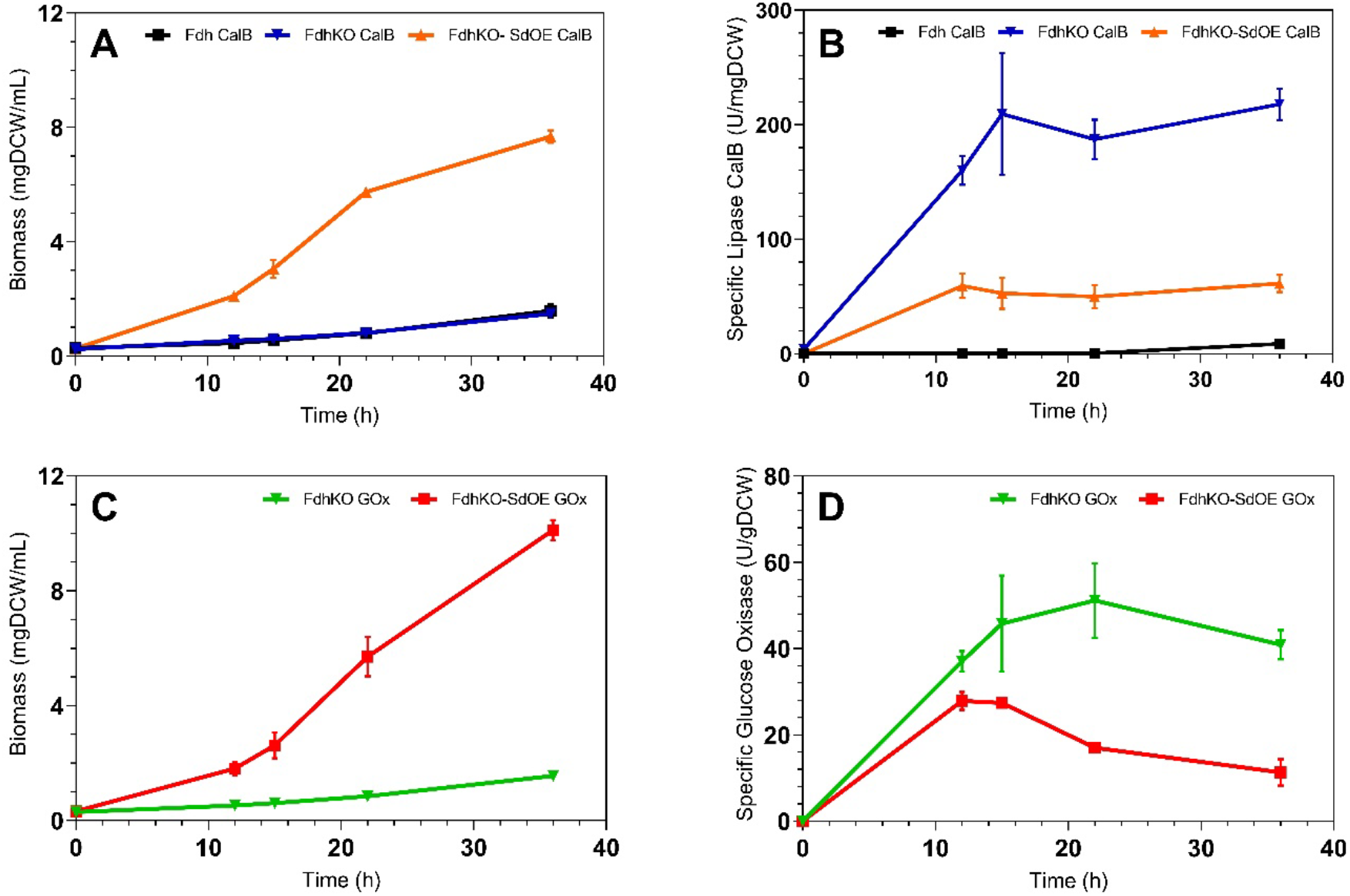
Biomass (panel A and C) and specific enzyme activity (panel B and D) during growth of FdhKO CalB (blue), FdhKO GOx (green), FdKO SdOE CalB (orange) and FdKO SdOE GOx (red) strains on YNB minimal medium containing sorbitol (YNBS). Data are the mean and standard deviation of triplicate cultures conducted in shake flasks. Lipase and glucose oxidase assays were performed in triplicates.

### Low oxygen transfer conditions favor the induction of pAOX1 of FdhKO strains grown on sorbitol-based methanol-free medium

Low oxygen availability has been reported to positively influence p*AOX1* induction for cells grown on methanol (Velastegui *et al*., 2023). This has also been reported for Fab-producing strains grown under carbon-limited conditions at different oxygen supplies (Carnicer *et al*., 2009). To examine a similar effect on sorbitol media, FdhKO eGfp and FdhKO SdOE eGfp strains were cultivated under different oxygen transfer conditions (OTC). This was achieved by varying the ratio between volumes of the culture medium and the culture flask, yielding culture systems characterized by different volumetric oxygen transfer coefficients (K_L_a; Gorczyca et al., 2020). K_L_a values of 10, 50, and 100 h^−1^ were used as benchmarks for low, medium, and high OTC levels, respectively.

For the FdhKO eGfp strain, the value of biomass at 22 h was within the same range for all culture conditions tested (0.62 mgDCW mL^−1^ on average, Figure 3A). Conversely, significant differences were observed for the FdhKO SdOE eGfp strain, with biomass values of 2.4, 6.3, and 7.4 mgCDW mL^−1^ on average under low, medium, and high OTC, respectively. Therefore, oxygen transfer was most likely a limiting factor for the growth of the FdhKO SdOE eGfp strain compared to the FdhKO eGfp strain. Under all tested culture conditions, the higher biomass observed in the FdhKO SdOE eGfp strain compared to the FdhKO eGfp strain is due to the overexpression of *SOR1*, which allows the FdhKO SdOE eGfp strain to metabolize sorbitol at a higher rate. At the sampling time, cell growth was not limited by sorbitol availability, as it remained in excess in the culture medium (i.e., at 22h, Figure S7).

**Figure 3.**
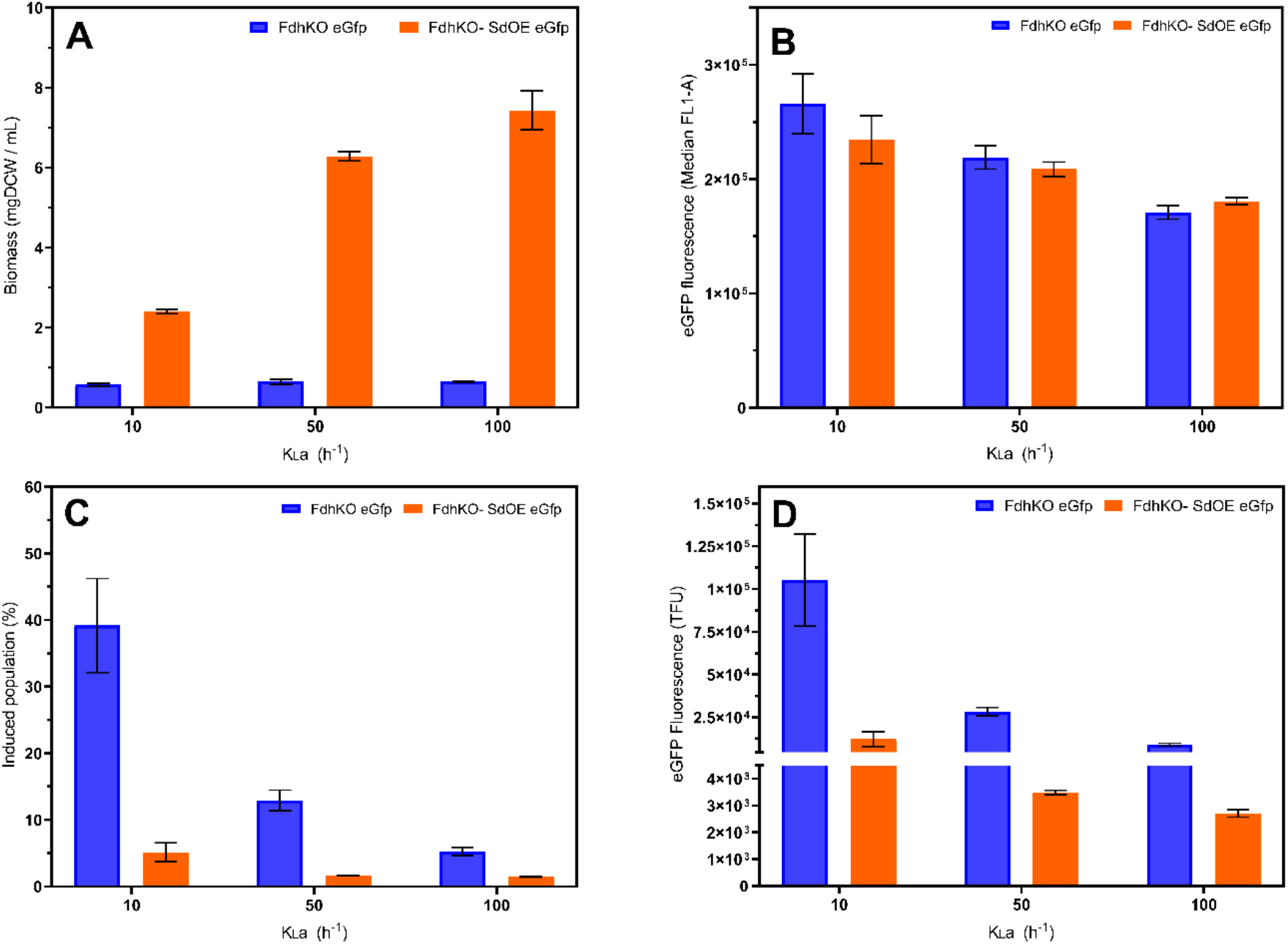
Biomass (panel A), Median FL1-A (panel B), induced population (panel C) and specific eGFP fluorescence (panel D) for FdhKO eGfp and FdhKO-SdOE eGfp strains cultured strains grown in YNBS2 under different OTC (low, K_L_a 10 h^-1^; medium, K_L_a 10 h^-1^; high, K_L_a 100 h^-1^), in shake-flask cultures. Culture samples were taken at the mid-exponential growth phase (i.e. after 22 h). eGFP fluorescence was quantified by flow cytometry on 20,000 cells and expressed as TFU (total fluorescence, see materials and method for calculation details). Data are means and standard deviations of triplicate cultures.

Cells were also analyzed by flow cytometry to assess the distribution of the cell population according to their eGFP fluorescence signal, distinguishing between non-induced and induced cells (*i*.*e*., with eGFP signal above a defined threshold, Theron et al., (2019)). The eGFP signal, which reflects the p*AOX1* induction level per cell decreased as the oxygen transfer increased (i.e., 36% for FdhKO eGfp and 23% for FdhKO SdOE eGfp strains, on average). It was not markedly different between the two strains for a given OTC (with a maximum of 12%) (Figure 3B). By contrast, a significant difference was observed in the fraction of the cell population in an induced state between the two strains and the culture conditions. The FdhKO SdOE eGfp strain exhibited a significantly lower fraction of the population in an induced state compared to the FdhKO eGfp strain, especially at low OTC, with a 7.8-fold difference (i.e., 5.1% vs. 39.2%, respectively; Figure 3C). The fraction of the population in an induced state also decreased significantly as oxygen transfer increased, with a 7.6-fold decrease for the FdhKO eGfp strain (i.e., 39.2% vs. 5.2%, respectively). Therefore, at low oxygen transfer, a greater portion of cells are in an induced state. This observation aligns with our previous findings of elevated *AOX1* expression levels under transient hypoxic conditions (Velastegui *et al*., 2023). In experiments using a p*AOX1*-*eGFP* strain grown on methanol, phenotypic diversification toward a highly fluorescent phenotype was observed under transient anoxic stress, with the fraction of the population in the highly fluorescent state increasing as hypoxic stress intensified.

By combining the eGFP fluorescence and the percentage of induced cells, the total fluorescence was calculated. It increased 12-fold for the FdhKO eGfp strain at low OTC compared to high one (i.e., 105,267 and 8,939 TFU, respectively). A similar trend was observed for the FdhKO SdOE eGfp strain, but to a lesser extent, with a 5-fold decrease (i.e., 12,184 and 2,709 TFU, respectively). The total eGFP fluorescence signal, and thus p*AOX1* induction, was 39-fold higher for the FdhKO eGfp strain at low OTC compared to the FdhKO SdOE eGfp strain at high OTC (i.e., 105,267 and 2,709 TFU, respectively) (Figure 3D). At low OTC, the e*GFP* gene expression level for the FdhKO eGfp strain was 5.4-fold higher than for the FdhKO SdOE eGfp strain (Figure S8). According to these results, the fraction of the cell population in an induced state is greatly affected by the level of oxygen transfer as compared to the strength of induction. Strain behavior is also influenced by the cells ability to metabolize sorbitol (i.e., *SOR1* expression).

### Understanding the differences in pAOX1 induction levels between FdhKO and FdhKO-SdOE strains grown on sorbitol-based methanol-free medium

In yeast, cytoplasmic formate serves as an intermediate in the THF-C1 pathway, the primary source of C1 units for *de novo* purine synthesis and energy storage molecules (e.g., adenosine triphosphate, ATP; Christensen and Mackenzie, 2006, Figure S9). Disruption of the Fdh encoding gene has been identified as a key factor in intracellular formate accumulation in cells grown on sorbitol, thereby enhancing p*AOX1* induction in a sorbitol-based methanol-free medium (Bustos *et al*., 2024). However, the reduced eGFP-specific fluorescence, lipase CalB and GOx activities observed in FdhKO-SdOE strains compared to their FdhKO counterparts (Figures 1 and 2) suggest that additional factors may influence p*AOX1* induction.

Overexpression of *SOR1* in FdhKO strains led to an increased sorbitol uptake rate, resulting in a higher growth rate (Table 2). This increased demand for anabolic precursors, including C1 units, likely reduces formate accumulation. To test this hypothesis, formate levels were quantified in the culture supernatants of FdhKO eGfp and FdhKO SdOE eGfp strains grown on sorbitol at different OTC. For the FdhKO SdOE eGfp strain, formate concentrations were markedly lower than for the FdhKO-eGfp strain across all tested OTC (Figure 4A). Additionally, formate content decreased for both strains as the oxygen transfer increased. For FdhKO eGfp strain, formate concentration was 32.3 µmol mgDCW^-1^ at low OTC, whereas it was nearly undetectable for the FdhKO SdOE eGfp strain at high OTC (0.33 µmol mgDCW^-1^). Sorbitol metabolism differs between FdhKO-eGfp and FdhKO-SdOE-eGfp strains due to *SOR1* overexpression. The FdhKO eGfp strain exhibited a low conversion efficiency of sorbitol into biomass, with a substrate-to-biomass yield coefficient (Y_x/s_) ranging from 0.18 to 0.22 g g^-1^ across all tested OTC. At low OTC, the Y_x/s_ of FdhKO SdOE eGfp strain was 1.9-fold higher than FdhKO eGfp (0.36 g g^-1^), increasing further to 0.7 g g^-1^ at medium and high OTC (Figure 4B). Conversely, when considering eGFP as a product, the substrate-to-product yield coefficient (Y_p/s_) for the FdhKO SdOE eGfp strain was 18-fold lower than that of the FdhKO eGfp strain at low OTC (38,586 vs. 2,110 TFU g^-1^, respectively, Figure 4C). For both strains, Y_p/s_ decreased as oxygen transfer increased, with a 10.9-fold reduction between low and high OTC for the FdhKO eGfp strain (38,586 vs. 3,521 TFU g^-1^, respectively).

**Figure 4:**
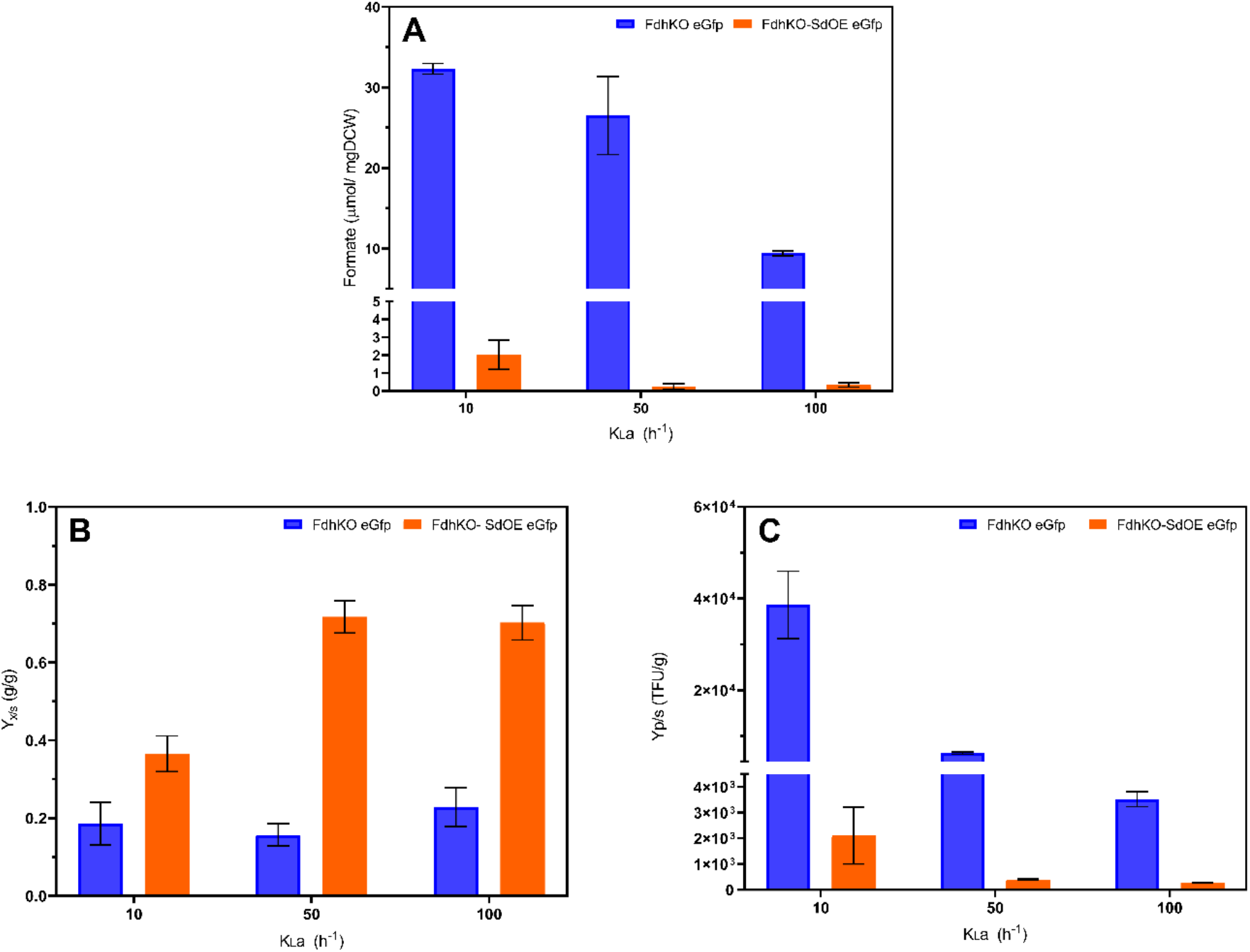
Specific formate concentration (panel A), biomass-substrate yield coefficient (Y_x/s_, panel B) and product-to-substrate yield coefficient (Y_p/s_, panel C) for FdhKO eGfp (blue bars) and FdhKO-SdOE eGfp (orange bars) strains grown under different OTC, benchmarked by K_L_a, in shake-flask cultures. Culture samples were taken at the mid-exponential growth phase (i.e. after 22 h). Data are means and standard deviations of biological triplicate.

Elevated formate concentrations were associated with low OTC, low Y_x/s_, and high Y_p/s_. Under these conditions, e*GFP* gene expression for the FdhKO eGfp strain was 5.4-fold higher than under conditions of high Y_x/s_ and low Y_p/s_ (i.e., high OTC, Figure S8). This suggests that when less substrate energy is directed toward biomass formation and instead allocated to secondary metabolism (e.g., eGfp synthesis), fewer C1 units are used in anabolic pathways, resulting in greater formate accumulation. This observation highlights the role of low Y_x/s_, favored under low OTC, in enhancing p*AOX1* induction.

## Conclusion

We recently demonstrated that *Komagataella phaffii* FdhKO strains exhibit self-induction of the p*AOX1*-based expression system when grown on sorbitol. In this study, we further show that these strains undergo diauxic growth in methanol-free glycerol-sorbitol media, effectively decoupling cell growth from recombinant protein production. Under these conditions, accumulated formate from the THF-C1 metabolism triggers p*AOX1* induction. To increase energy generation in FdhKO strain, sorbitol catabolism was optimized by overexpression of *SOR1*. Although increased sorbitol metabolism enhances biomass formation, it simultaneously reduced p*AOX1* induction, as evidenced by lower formate accumulation, decreased eGFP fluorescence, and reduced secreted CalB and Gox activities in *SOR1*-overexpressing strains.

Oxygen transfer also modulates these dynamics, with lower oxygen availability favoring higher p*AOX1* induction due to increased formate accumulation. Notably, the observed increase in p*AOX1* expression appears to result from phenotypic diversification within the cell population, leading to a higher fraction of induced cells rather than an overall increase in p*AOX1* induction per cell. These findings underscore the metabolic trade-offs in in sorbitol-based processes and highlight the potential for optimizing carbon flux to balance recombinant protein production and cell growth. Future studies should focus on fine-tuning metabolic pathways to enhance energy efficiency while sustaining robust p*AOX1* induction in methanol-free, sorbitol-based media.

## AUTHOR CONTRIBUTIONS

**Cristina Bustos and Rocio Cozmar:** Conceptualization; data curation; formal analysis; investigation; methodology; validation; visualization; writing – original draft; writing-review and editing. **Patrick Fickers:** Conceptualization; formal analysis; investigation; methodology validation; validation; visualization; funding acquisition; resources; supervision; writing – original draft; writing-review and editing. **Julio Berrios:** funding acquisition; writing-review.

### ACKNOWLEDGEMENTS

The authors thank L. Henrion, V. Vandenbroucke, A. Padmanabhan, Korka, S. Telek, A. Zicler, S. Steels, and R. Thomas for their technical assistance and valuable discussions. Graphical abstract was created with BioRender.com.

## DATA AVAILABILITY

Data are available upon request to the corresponding author

