## Supplementary figures and tables for "Sorbitol uptake and oxygen transfer shape *AOX1* promoter induction under methanol-free conditions in *Komagataella phaffii* lacking formate dehydrogenase"

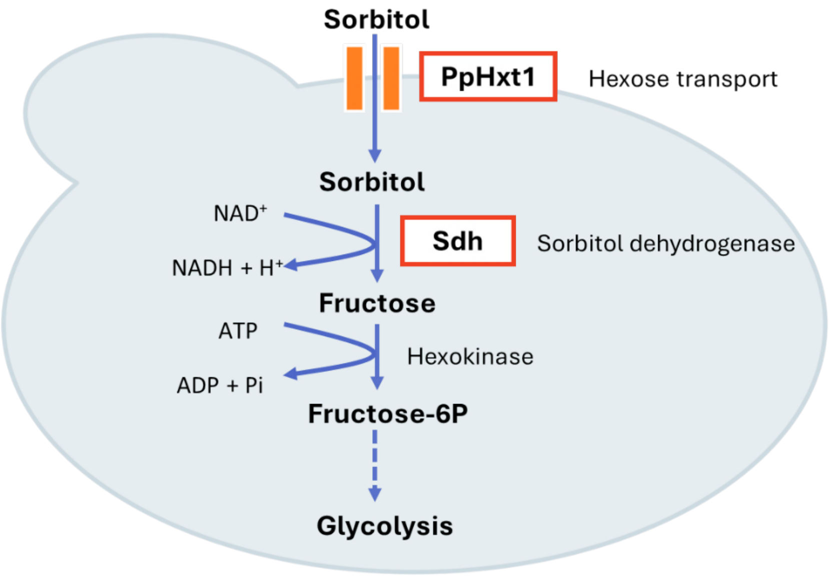


**Figure S1.** Sorbitol metabolism in *K. phaffii*: PpHxt1, a hexose transporter, is evaluated in this investigation as a putative sorbitol transporter; Sdh refers to sorbitol dehydrogenase.

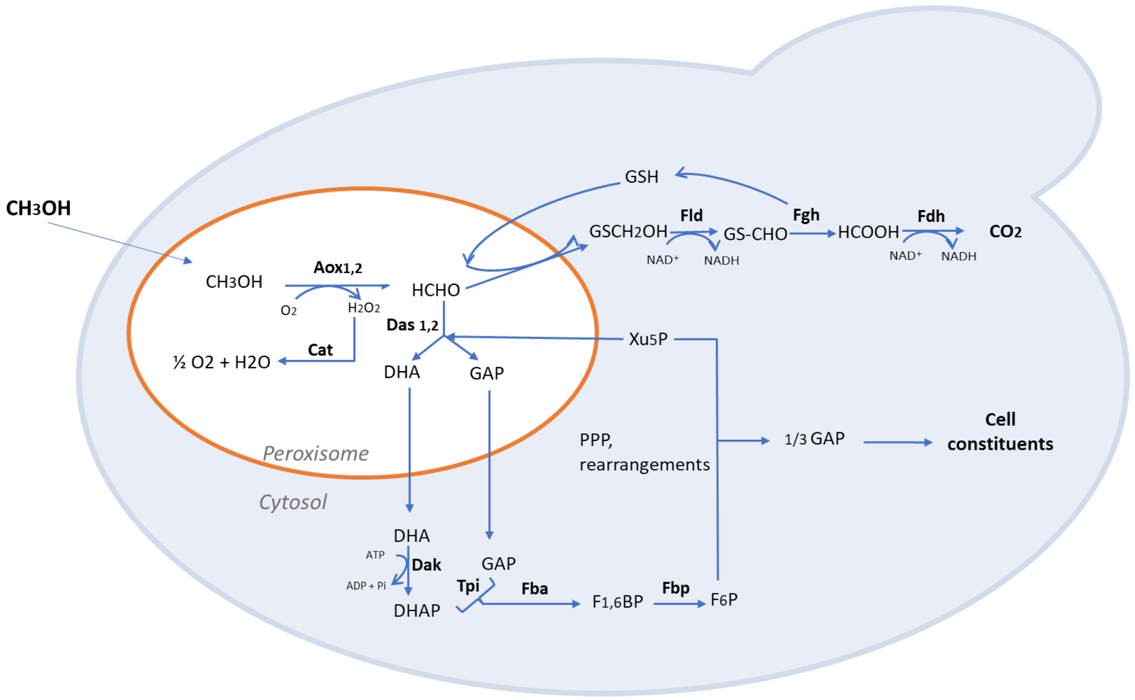


**Figure S2**. Methanol utilization pathway in *Komagataella phaffii*. Aox1, 2: alcohol oxidase1 and 2; Cat: catalase; Das1, 2: dihydroxyacetone synthase; Fld: formaldehyde dehydrogenase; Fgh: S-formylglutathione hydrolase; Fdh: formate dehydrogenase; Dak: dihydroxyacetone kinase; Tpi: triose phosphate isomerase; Fba: fructose-1,6-bisphosphate aldolase; Fbp: fructose-1,6-bisphosphatase; GS(H): glutathione; DHA: dihydroxyacetone; DHAP: dihydroxyacetone phosphate; GAP: glyceraldehyde-3-phosphate; F1,6BP: fructose-1,6-bisphosphate; F6P: fructose-6-phosphate; Xu5P: xylulose 5-phosphate.

```
XP_002489933 - K. phaffii GS115   1    MS--DNPSVILKRINEIVIEDRPIPAIEDPHYVKIAIKKTGICGSDVHFYTDGCCGSFKL  58
                                       MS   NP+V+L+++ +I IE RPIP I+DPHYVK+AIK TGICGSD+H+Y  G  G + L
XKU26523.1 - S. cerevisiae        1    MSQNSNPAVVLEKVGDIAIEQRPIPTIKDPHYVKLAIKATGICGSDIHYYRSGGIGKYIL  60

XP_002489933 - K. phaffii GS115   59   ESPMVLGHESAGIVVEVGSEVKSLRVGDKVACEPGIPSRYSNAYKSGHYNLCPEMAFAAT  118
                                       ++PMVLGHES+G VVEVG  V  ++VGD+VA EPG+PSRYS+  K G YNLCP MAFAAT
XKU26523.1 - S. cerevisiae        61   KAPMVLGHESSGQVVEVGDAVTRVKVGDRVAIEPGVPSRYSDETKEGRYNLCPHMAFAAT  120

XP_002489933 - K. phaffii GS115   119  PPIDGTLCRYFLLPEDFCVKLPEHVSLEEGALVEPLSVAVHAARLAKITFGDSVVVFGAG  178
                                       PPIDGTL +Y+L PEDF VKLPE VS EEGA VEPLSV VH+ +LA + FG  VVVFGAG
XKU26523.1 - S. cerevisiae        121  PPIDGTLVKYYLSPEDFLVKLPEGVSYEEGACVEPLSVGVHSNKLAGVRFGTKVVVFGAG  180

XP_002489933 - K. phaffii GS115   179  PVGLLVAATARAYGATNVLIVDIFDDKLTLAKDTLQVATHSFNS--------KNGMDNLL  230
                                       PVGLL  A ARA+GAT+V+ VD+FD+KL  AKD    AT++FNS        ++  D +
XKU26523.1 - S. cerevisiae        181  PVGLLTGAVARAFGATDVIFVDVFDNKLQRAKDF--GATNTFNSSQFSTDKAQDLADGVQ  238

XP_002489933 - K. phaffii GS115   231  ESFEGKHPNVSIDCTGVESCIAAGINALAPRGVHVQVGMGKSEYNNFPLGLICEKECIVK  290
                                       +   G H +V  +C+G + CI A +      G  VQVGMGK+ Y NFP+  +  KE   +
XKU26523.1 - S. cerevisiae        239  KLLGGNHADVVFECSGADVCIDAAVKTTKVGGTMVQVGMGKN-YTNFPIAEVSGKEMKLI  297

XP_002489933 - K. phaffii GS115   291  GVFRYCYNDYNLAVELIASGKVEVKGLVTHRFKFTEAVDAYD--TVRQGKAIKAIIDGPE  348
                                       G FRY + DY  AV L+A+GKV VK L+TH+FKF +A  AYD      G+ +K II GPE
XKU26523.1 - S. cerevisiae        298  GCFRYSFGDYRDAVNLVATGKVNVKPLITHKFKFEDAAKAYDYNIAHGGEVVKTIIFGPE  357
```

**Figure S3**. Sorbitol Dehydrogenase enzyme. Protein alignment. XP_002489933.1 corresponds to (gene PAS_chr1-1_0490) protein sequence *K. phaffii*, and XKU26523.1 corresponds to *Saccharomyces cerevisiae*.

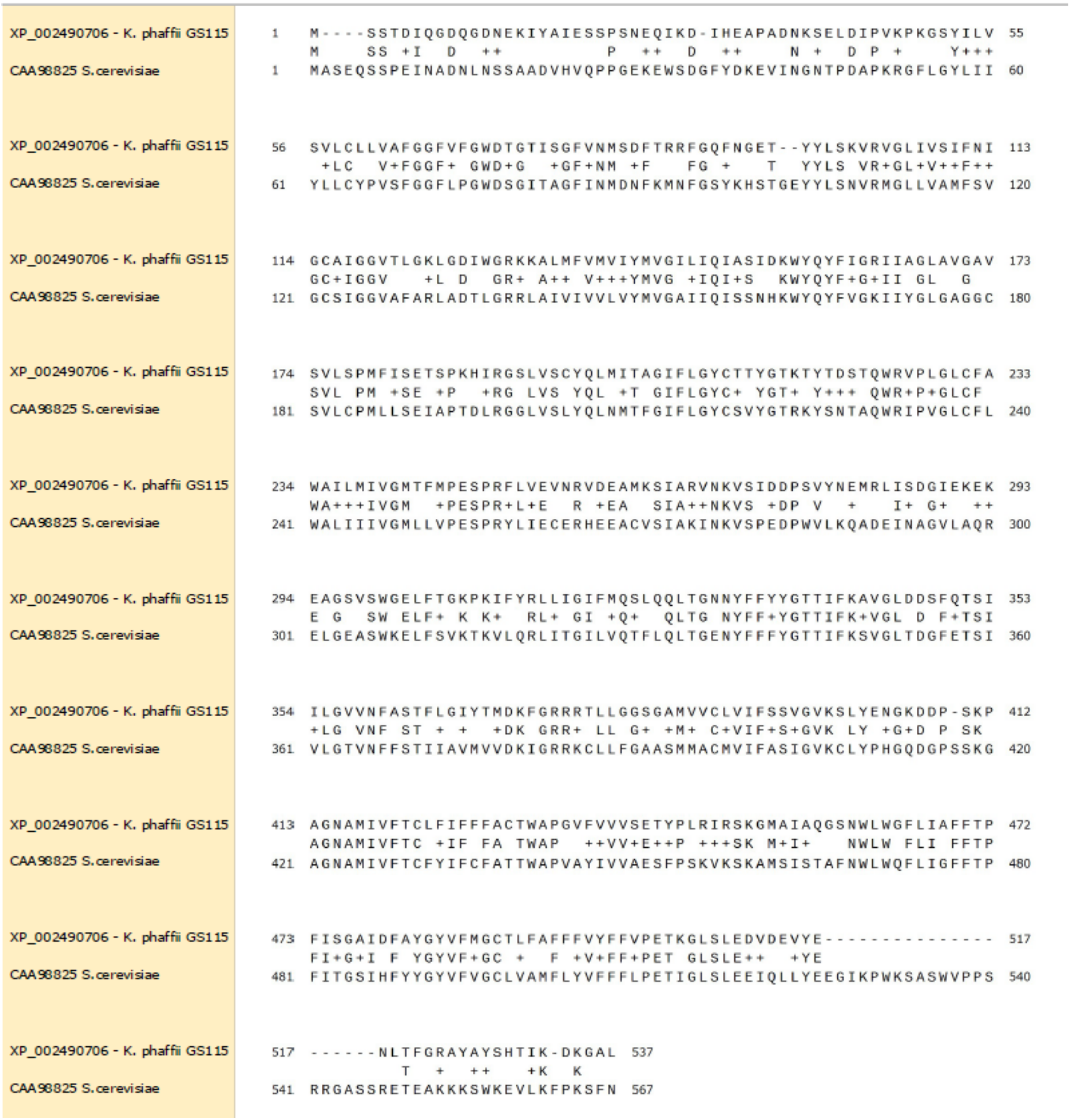


**Figure S4.** Hexose transporter protein. Protein alignment. XP_002490706.1 corresponds to (gene PAS_chr1-4_0570) protein sequence *K. phaffii*, and CAA98825.1 corresponds to *Saccharomyces cerevisiae*.

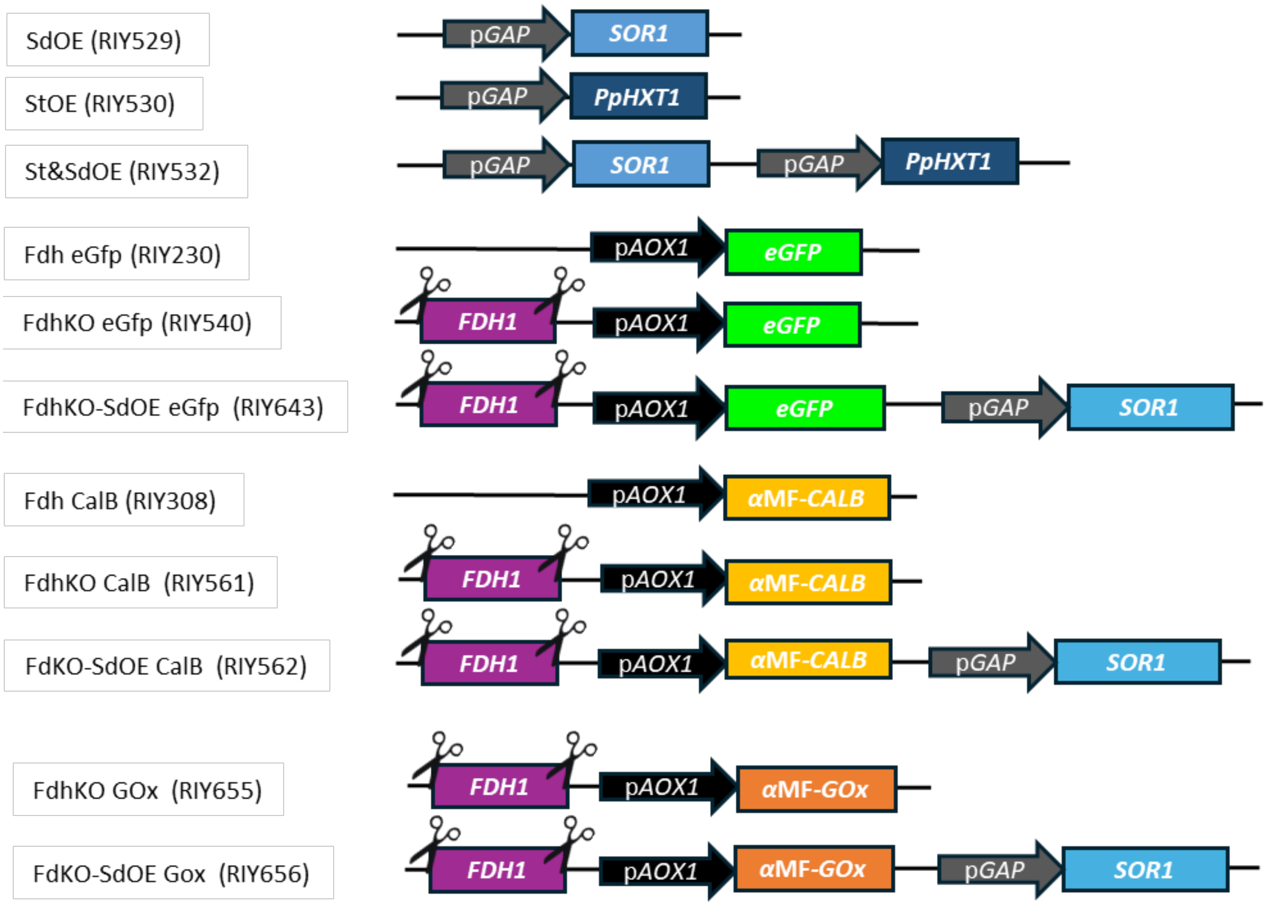


**Figure S5.** Schematic representation of the genotype of yeast strains.

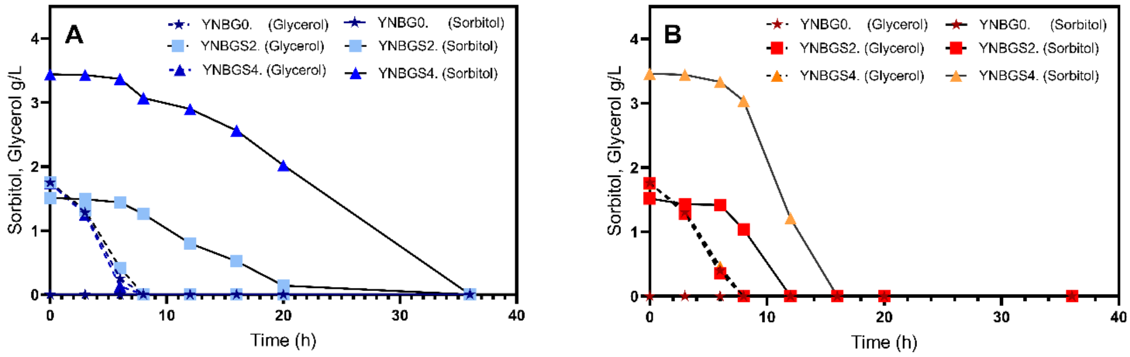


**Figure S6.** Glycerol and sorbitol concentration during the growth of FdhKO eGfp strain (blue colors, Panel A) and FdhKO SdOE eGfp strain (orange colors, Panel B) in YNB minimal medium containing different mixtures of glycerol and sorbitol (YNBG0, 1.75 g $L^{-1}$ glycerol; YNBGS2, 1.75 g $L^{-1}$ glycerol and 1.75 g $L^{-1}$ sorbitol; YNBGS4, 1.75 g $L^{-1}$ glycerol and 3.5 $L^{-1}$ sorbitol). The data show one representative culture.

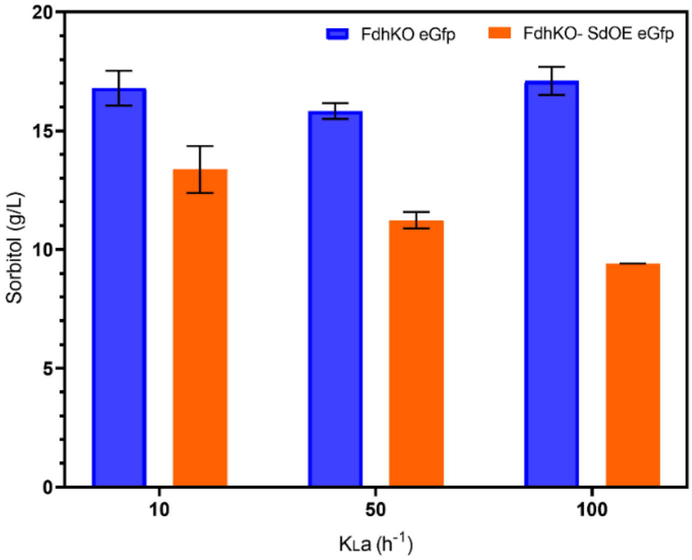


**Fig S7.** Sorbitol concentration after 22 h of growth of strains SdhKO eGfp (blue) and SdhKO SdOE eGfp (orange) in the YNBS2 medium under different OTC (low, $K_La$ 10 $h^{-1}$; medium, $K_La$ 10 $h^{-1}$; high, $K_La$ 100 $h^{-1}$). Data are the mean and standard deviation of triplicate cultures conducted in in shake flasks.

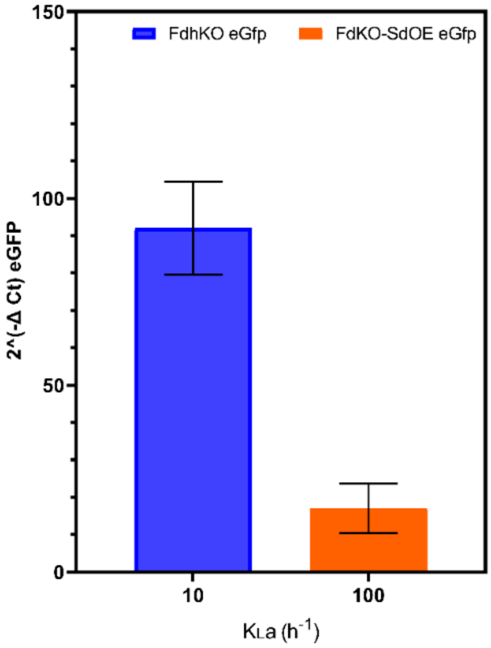


**Figure S8.** e*GFP* gene expression level for FdhKO eGfp and FdhKO-SdOE eGfp strains cultured strains grown under different OTC (low, $K_La$ 10 $h^{-1}$; medium, $K_La$ 10 $h^{-1}$; high, $K_La$ 100 $h^{-1}$), in shake-flask cultures. Culture samples were taken at the mid-exponential growth phase (i.e. after 22 h). Data are means and standard deviations of triplicate cultures. Gene expression levels were normalized according to that of actin gene.

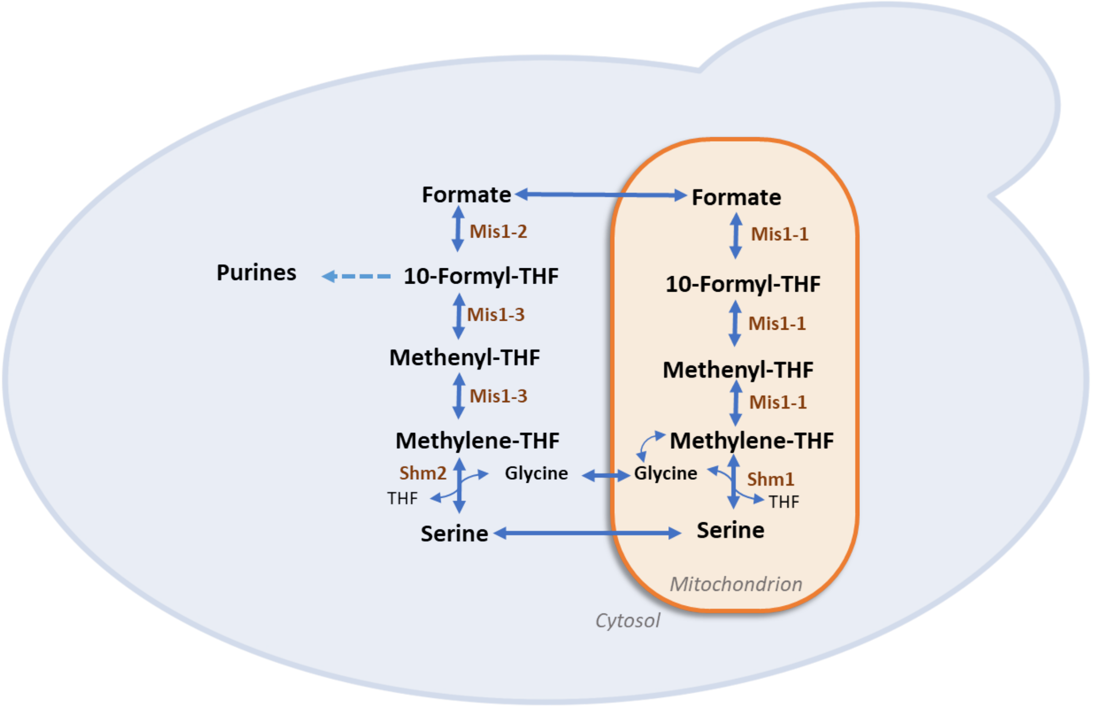


**Figure S9**. Tetrahydrofolate (THF) mediated one-carbon (THF-C1) metabolism in yeast. Fdh, formate dehydrogenase; Shm1, mitochondrial serine hydroxymethyltransferase; Shm2, cytosolic serine hydroxymethyltransferase; Mis1-2, formate-tetrahydrofolate ligase; Mis1-3, methylenetetrahydrofolate dehydrogenase & methenyltetrahydrofolate cyclohydrolase; Mis1-1, trifunctional enzyme: formate-tetrahydrofolate ligase, methenyltetrahydrofolate cyclohydrolase and methylenetetrahydrofolate reductase; THF, tetrahydrofolate. Figure adapted from (Bustos, et al.2024, Christensen, et al.2006)

**Table S1.** *Escherichia coli* strains used in this study.

| Strain name (plasmid) | Plasmid - genotype | Source/Reference |
|---|---|---|
| A2 | BB1_23 | (Prielhofer *et al.*, 2017) |
| D12 | BB3aZ_14 | (Prielhofer *et al.*, 2017) |
| A4 | BB1_12_p*GAP* | (Prielhofer *et al.*, 2017) |
| C1 | BB1_34_*ScCYC1*tt | (Prielhofer *et al.*, 2017) |
| C9 | BB1_34_*RPS3*tt | (Prielhofer *et al.*, 2017) |
| C12 | BB3aK_AC | (Prielhofer *et al.*, 2017) |
| D4 | BB2_AB | (Prielhofer *et al.*, 2017) |
| D5 | BB2_BC | (Prielhofer *et al.*, 2017) |
| RIE341 (RIP341) | A2_BB1_23_*SOR1* | This work |
| RIE343 (RIP343) | A2_BB1_23_Pp*HXT1* | This work |
| RIE354 (RIP354) | BB3-p*GAP*- Pp*HXT1-SsCy1*tt | This work |
| RIE355 (RIP355) | BB3-p*GAP-SOR1-SsCy1*tt | This work |
| RIE370 (RIP370) | BB3-AC-LoxP-Zeo-loxP | This work |
| RIE372 (RIP372) | BB2(AB)-p*GAP-SOR1- SsCy1*tt | This work |
| RIE374 (RIP374) | BB2(BC1)-P*GAP*-Pp*HXT1-RPS3*tt | This work |
| RIE378 (RIP378) | BB3a-p*GAP-SOR1-SsCy1*tt, p*GAP*- Pp*HXT1-RPS3*tt | This work |

**Table S2**. Primers used in this study

| Name | Sequence 5´to 3´ | Restriction site/gene |
|---|---|---|
| M13-Fw | GTAAAACGACGGCCAGT | |
| M13-RV | AACAGCTATGACCATG | |
| pGAP.Int-Fw | CGTCGCTGGCAATAATAGCGG | |
| *ScCYC1*tt.Int-Rv | GGGACCTAGACTTCAGGTTGTC | |
| Rps3tt.Int-Rv | GACGAGTCCAGGGCTATCTTAAG | |
| SOR1-Fw | C**GGTCTC**ACATGTCCGATAACCCAAGTGTTATCTTAA AAGGATTAATGAGATTGTCATAGAAGATAGACCAAT TCCAGCCATTGAGGATCCTCACTATGTGAAAATAGC AATCAAAAGACCGGAATTTG | BsaI |
| SOR1-Rv | C**GGTCTC**CAAGCTTTACTCTGGGCCGTCAATGATAG | BsaI |
| qAct-F | AGATGGCTCCGAGAAGTTCA | *ACT1* |
| qAct-R | GTTGCTCAGAGGGCTTCAAC | *ACT1* |
| qeGFP-F | ATCATGGCCGACAAGCAGAA | e*GFP* |
| qeGFP-R | TCTCGTTGGGGTCTTTGCTC | e*GFP* |
| ST_qPCR_Fo | CCAGGTGTTTTCGTCGTTGT | Pp*Hxt1* |
| ST_qPCR_Rev | AGGCGAACAGAGTACATCCC | Pp*Hxt1* |
| SDH_qPCR_Fo | CCCGTCTCGTTACAGCAATG | *SOR1* |
| SDH_qPCR_Rev | GCATGGACAGCAACACTCAA | *SOR1* |
